# Reply to “A discriminative learning approach to differential expression analysis for single-cell RNA-seq”

**DOI:** 10.1101/648733

**Authors:** Etienne Becht, Edward Zhao, Robert Amezquita, Raphael Gottardo

**Affiliations:** Vaccine and Infectious Disease Division, Fred Hutchinson Cancer Research Center, Seattle, WA, 98109, USA; Department of Biostatistics, University of Washington, Seattle, WA, 98195, USA

## Abstract

Multivariate logistic regression (mLR) has been recently proposed by Ntranos et al. to perform gene differential expression analyses of single-cell RNA-sequencing (scRNAseq) data. Herein we reproduce and extend some of their findings. We notably show that while mLR performs better in simulated datasets, these simulations do not recapitulate important features of experimental datasets. Indeed, our results suggest that MAST followed by Sidak aggregation of the p-values perform better than mLR on experimental datasets. Overall, we highlight that most of the new results obtained by Ntranos et al is likely due to the quantification of scRNAseq data at the transcript or transcript compatibility classes level, rather than the use of mLR.

## Main text

In *A discriminative learning approach to differential expression analysis for single-cell RNA-seq*^*1*^, Ntranos, Yi, Melsted and Pachter propose multivariate logistic regression (mLR) as a way to perform gene differential expression (GDE) on sub-gene features for single cell RNA-sequencing (scRNAseq) data. While in our hands mLR performs better than other methods in simulated datasets, these simulations do not recapitulate important features of experimental datasets. Indeed, our results suggest that MAST followed by Šidák aggregation of the p-values performs better than mLR on experimental datasets. Overall, we highlight that most of the new results obtained by Ntranos et al. are likely due to the quantification of scRNAseq data at the transcript or transcript compatibility classes level, rather than the use of mLR.

Ntranos et al. restate GDE as a classification problem. Specifically, their method models the probability of a cell belonging to one of two groups (for instance cell types, clusters or treatment groups) by using the count matrix of transcripts or transcript compatibility classes (TCCs) mapping to a gene *g* as predictor variables. Likelihood-ratio tests are subsequently used to compare the full model with all TCCs as predictors to a null model with a simple intercept, providing a p-value for gene *g*. The method relies on two main ideas, namely 1) the use of logistic regression as a way of performing gene differential expression from single-cell RNAseq data quantified at the transcript or TCC level, and 2) the usefulness of RNAseq quantification at the transcript or TCC level to perform GDE analysis. Both of these aspects are interesting to study but may independently provide benefits. Their respective contribution to the results presented should therefore be separately evaluated. In this reply, we argue that the majority of the results presented in the original paper should be attributed to the aggregation of transcript-level expression to the gene level, while multivariate LR itself yielded contrasting results in our hands. In particular, we show that using p-value aggregation method to obtain gene-level p-values from data summarized at the sub-gene level, a topic which some of the same authors published about^2^, also improves the performance of other differential expression methods. We notably highlight that using Model-based Analysis of Single-cell Transcriptomics (MAST)^3^ on sub-gene features in conjunction with Šidák aggregation of the p-values to the gene level seems to perform better than mLR on real (non-simulated) datasets.

By analyzing the sorted human T cell data^4^ shared by Ntranos et al, we observed, consistent with their findings, that mLR performed better than applying LR on each TCC followed by p-value aggregation using Bonferonni correction or LR performed on data quantified at the gene level (Figures 1a, 1b, 1c). However, we found that the MAST method^3^ performed at the TCC level followed by p-value aggregation using the Šidák method (MAST-Šidák)^5^ also identified the CD45 gene (PTPRC) as differentially-expressed when comparing naive and memory CD4 T cells (Figure 1a) or naive CD8 and memory CD4 T cells (Figure 1b) but not naive CD4 and memory CD8 T cells (Figure 1c). In this particular example, the observed results are thus likely better explained by the level at which the data has been quantified (using TCCs) rather than the LR method itself. Performing differential expression analysis with MAST at the TCC level followed by aggregation of the p-values to the gene level (as promoted by some of the same authors in another publication^2^) yields results that are consistent with prior immunological knowledge and that lead to the same conclusions as results obtained using mLR.

**Figure 1.**
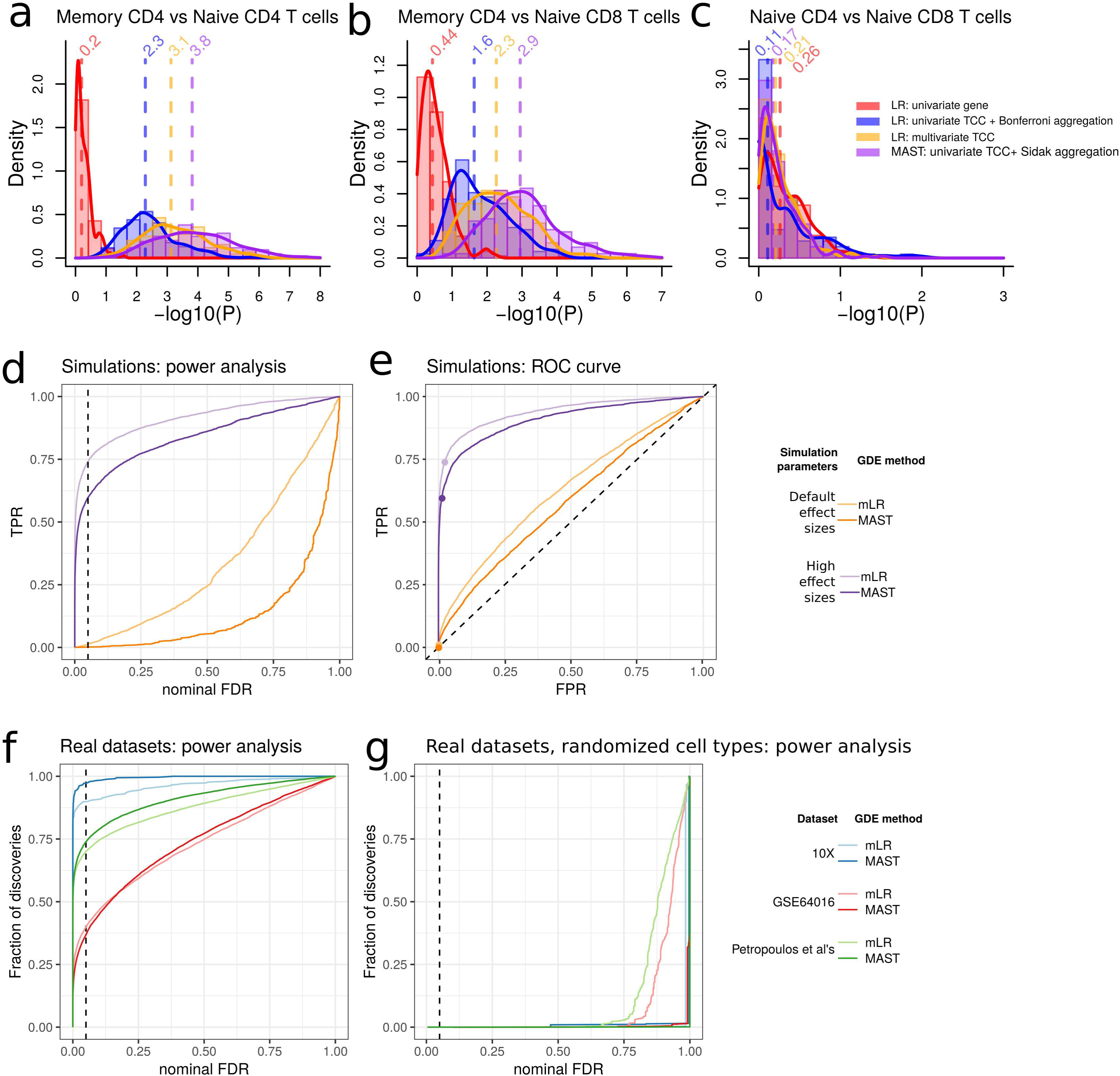
Distributions of -log10(p-values) generated by logistic regression on gene counts (red), univariate logistic regression on TCCs followed by Bonferronni aggregation of p-values at the gene level (blue), multivariate logistic regression (mLR) on TCCs (yellow) or univariate MAST on TCCs followed by Šidák aggregation of the p-values (purple), for 200 subsamples of 2,000 cells out of 3,000 **a)** memory or naive CD4 T cells, **b)** memory CD4 or naive CD8 T cells and **c)** naive CD4 or naive CD8 T cells. In each panel, vertical dashed lines indicate the median - log10(p value) of the 200 random subsamples. **d**) power and **e**) ROC curve analyses for two simulations (default or high effect sizes for differential expression) and two gene differential expression methods (mLR and MAST-Šidák). **f**) power analysis and **g**) power analysis with randomized cell type labels in three real datasets.

The results presented in Supplementary Figure 6a of Ntranos et al. are used to support the uniqueness of the genes identified by mLR. In this figure, the authors selected the 3,000 genes with the lowest p-values for each method. However, the authors used Seurat^6^ to apply MAST^3^, Monocle - Tobit^7^ and DESeq2^8^. The Seurat package applied two filters to the gene matrices (a 10% expression frequency filter and a 0.25 minimum fold change filter) resulting in more than 50% of the genes being unassayed by these three methods (Supplementary Figure 1A). Subsetting on the 11,688 genes assayed by all methods instead showed that for any method, 1,347 out of the top 3,000 genes are common to all five methods (Supplementary Figure 1B), rather suggesting a strong consensus among differential expression methods on this dataset.

In the analysis of simulated data, it appears that the authors performed univariate analyses at the gene level for all methods, only performing aggregation of sub-gene features to the gene level for logistic regression. However, in these simulations, MAST also benefits from p-value aggregation, which increases its true positive rate (TPR) without increasing its false positive rate (FPR) (Supplementary Figure 2). This analysis also revealed that in the original simulated dataset, both methods only reached a low detection rate at a false discovery rate (FDR) threshold of 0.05 (Figure 1d). In other words, the effect size was so small that all methods would fail to detect most of the differentially expressed genes. Increasing the fold change for differentially-expressed genes in this dataset simulated with the splatter package^9^ greatly increased the detection rate, from 0.0014% to 60% and from 0.013% to 74% of the truly differentially-expressed genes being detected by MAST-Šidák and mLR, respectively (Figure 3d). In any case, these results showed that mLR has higher TPR and FPR at any FDR cut-off than MAST in the simulated datasets. These simulated results imply that MAST detects less genes as differentially-expressed compared to mLR; it is however unclear whether these apply to real experimental datasets.

The original manuscript does not present comparisons of mLR to other GDE techniques on non-simulated data. We benchmarked mLR and MAST-Šidák on three real datasets. Namely, we re-analyzed two datasets used in the original manuscript (the embryonic development Petropoulos et al. dataset (EMTAB3929^10^), the FACS-sorted CD4 naive and CD4 memory T cell dataset from Zheng et al.^4^ as well as the cell-cycle dataset from Leng et al. (GSE64016^11^). On each of these datasets and across all p-value cutoffs, MAST-Šidák had higher or similar frequencies of genes called as differentially-expressed compared to mLR (Figure 1f). The analysis of experimental datasets therefore contradicts the analysis of splatter-simulated datasets. While we were not able to identify the reason for this discrepancy, we noted several differences between this simulation framework and real datasets. Notably, the grouping of simulated transcripts into genes in simulated data is arbitrary. Therefore, the distribution of pairwise correlation coefficients for transcripts mapping to the same gene is similar to the one of transcripts mapping to different genes in the simulations. In contrast, in real datasets the distribution of correlation coefficients for transcripts mapping to the same genes is more right-skewed than the one of transcripts mapping to different genes (Supplementary Figure 3). It should be noted that while Bonferroni aggregation of p-values is more conservative than Šidák’s, it does not assume independence of the tests being combined (i.e. transcripts). However, this assumption is unlikely to be an issue here given that most transcripts are weakly correlated (Supplementary Figure 3). When transcripts are in fact correlated, our data shows that the dependence is positive, in which case Šidák aggregation should be conservative^12^. While we cannot measure TPR and FPR directly on these real datasets, we performed GDE analyses with randomly shuffled cell type labels as a proxy for FPR. In this setting, both methods identified virtually no gene as differentially-expressed at a FDR cut-off of 0.05 (Figure 1g). These results suggest that the higher discovery rate of MAST in experimental datasets is unlikely to be caused by a substantially higher FPR.

In conclusion, we agree that using sub-gene features to perform gene expression analyses in scRNAseq experiments is useful and may reveal subtle differences across cell populations that would otherwise be left undetected. However, we question whether logistic regression is truly the best framework to do so, given using Šidák aggregation on MAST leads to results that are comparable if not better on empirical data. In addition, LR is more restrictive in terms of modeling as it is restricted to discrete outcomes and would make it more difficult to adjust for biological or technical covariates that directly affect gene expression (e.g. batch effects). Finally, it should be noted that we anticipate that our MAST results would likely extend to other univariate methods where Šidák aggregation can be applied. Herein, we focused on MAST for simplicity as we are familiar with the software and it has been shown to perform well in benchmarks^13^, but it is likely that other differential expression methods such as DeSeq2^8^, Monocle-Tobit^7,8^ and SCDE^14^ would also benefit from p-value aggregation in these benchmarks.

---

*Supplementary figure 1*

**a**) Number of features assayed (left) and reproduction of Supplementary Figure 6 of the original paper (right). The right panel represents the intersections of the gene sets composed for each method by the 3,000 genes with lowest p-values. **b**) Corresponding figure when genesets intersections are computed on the features assayed by all five differential expression methods.

*Supplementary figure 2*

Across all simulations (see Supplementary Table 1 for the corresponding parameters), biplots of the false positive rate (left) and true positive rate (right) for MAST performed on gene counts (x-axes) versus MAST with p-values aggregation of transcripts to genes (y-axes).

*Supplementary figure 3*

Log10 density estimates of the distribution of correlation coefficients for simulated (left) or experimental (right) datasets for transcripts mapping to the same gene (red) or random transcripts (blue). For splatter, we show results on datasets generated using three sets of parameters: i) as per Ntranos et al. (*authors*), ii) with high fold change (*authors_highES)* and iii) default parameters (*vanilla*).

## Abbreviations

FPR: false positive rate
GDE: gene differential expression
LR: logistic regression
MAST: Model-based Analysis of Single-cell Transcriptomics^3^
mLR: multivariate logistic regression
scRNAseq: single-cell RNA sequencing
TCC: transcript compatibility classes
TPR: true positive rate

## Supplementary methods

Most of the analysis presented here are extensions of the analysis performed in the original article. The original code is available at https://github.com/pachterlab/NYMP_2018. Specifically, Figure 1a, 1b and 1c are adapted from the code available at https://github.com/pachterlab/NYMP_2018/tree/master/10x_example-logR, while figures 1d, 1e, 1f, 1g and supplementary figures 2 and 3 are adapted from code available at https://github.com/pachterlab/NYMP_2018/tree/master/simulations. Supplementary figure 1 is adapted from code available at https://github.com/pachterlab/NYMP_2018/tree/master/embryo. The adapted code is available at https://github.com/ebecht/logistic_regresion_for_GDE

### Datasets

We analyzed three publicly-available scRNAseq datasets as well as datasets simulated using the R splatter package^9^. Public datasets originated from Petropoulos et al^10^ (available from ArrayExpress or conquer^13^, accession number EMTAB3929), Leng et al^11^ (available from Gene Expression Omnibus or conquer^13^, accession number GSE64016). These two datasets were analyzed using transcripts as features. The third empirical dataset originates from Zheng et al. and was quantified at the TCC level by Nnatros et al^1^. For each dataset, we filtered-out transcripts (or TCCs) with 0 detected counts in more than 20% of the cells, except for the analysis of CD45 (Figure 1a,b,c) where we used a 0.1% expression frequency threshold as in the original publication).

Simulated datasets were produced using the splatter R package (v1.6.1). We used splatter parameters as shared by Nnatros et al. with either default fold change parameter (0.1) or high fold change parameter (1.5).

#### Algorithms

For logistic regression we used the *glm* function of the *stats* R package (v3.5.1) and performed likelihood ratio tests using the lmtest R package (v0.9-36). We used MAST (v1.9.1) in conjunction with aggregation of the transcript or TCC-level p-values to gene p-values using the Šidák method^5^. Briefly, the Šidák method aggregates a vector p_i=1…n_ of p values to p_Šidák_=1-(1-min_1≤i≤n_(p_i_))^n^ where min(.) denotes the minimum. We used the first order approximation p_Šidák_=n*min_1≤i≤n_(p_i_) when very low p-values led to numerical errors. The Tobit, SCDE and DESeq2 models were used as per the original paper.

To estimate the mean correlation of transcripts mapping to a given gene, for each gene we computed the average of the upper triangle of the correlation matrix of the transcripts mapping to it. For randomly selected genes, we randomly selected 2,000 genes from each dataset and computed the same metric.

