## Supplementary figures and images for "Reply to “A discriminative learning approach to differential expression analysis for single-cell RNA-seq”"

### Supplementary Figure 1

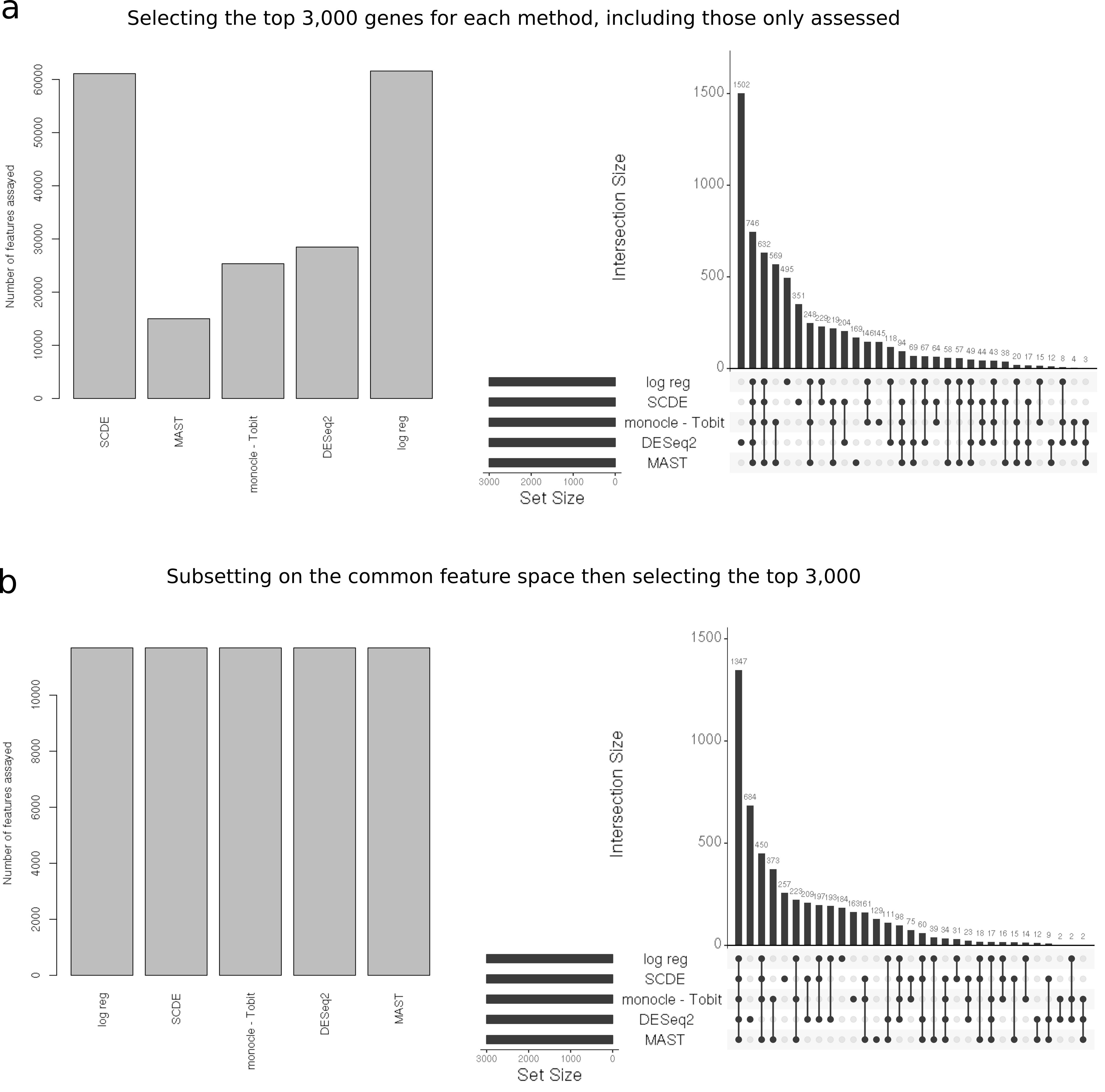

### Supplementary Figure 2

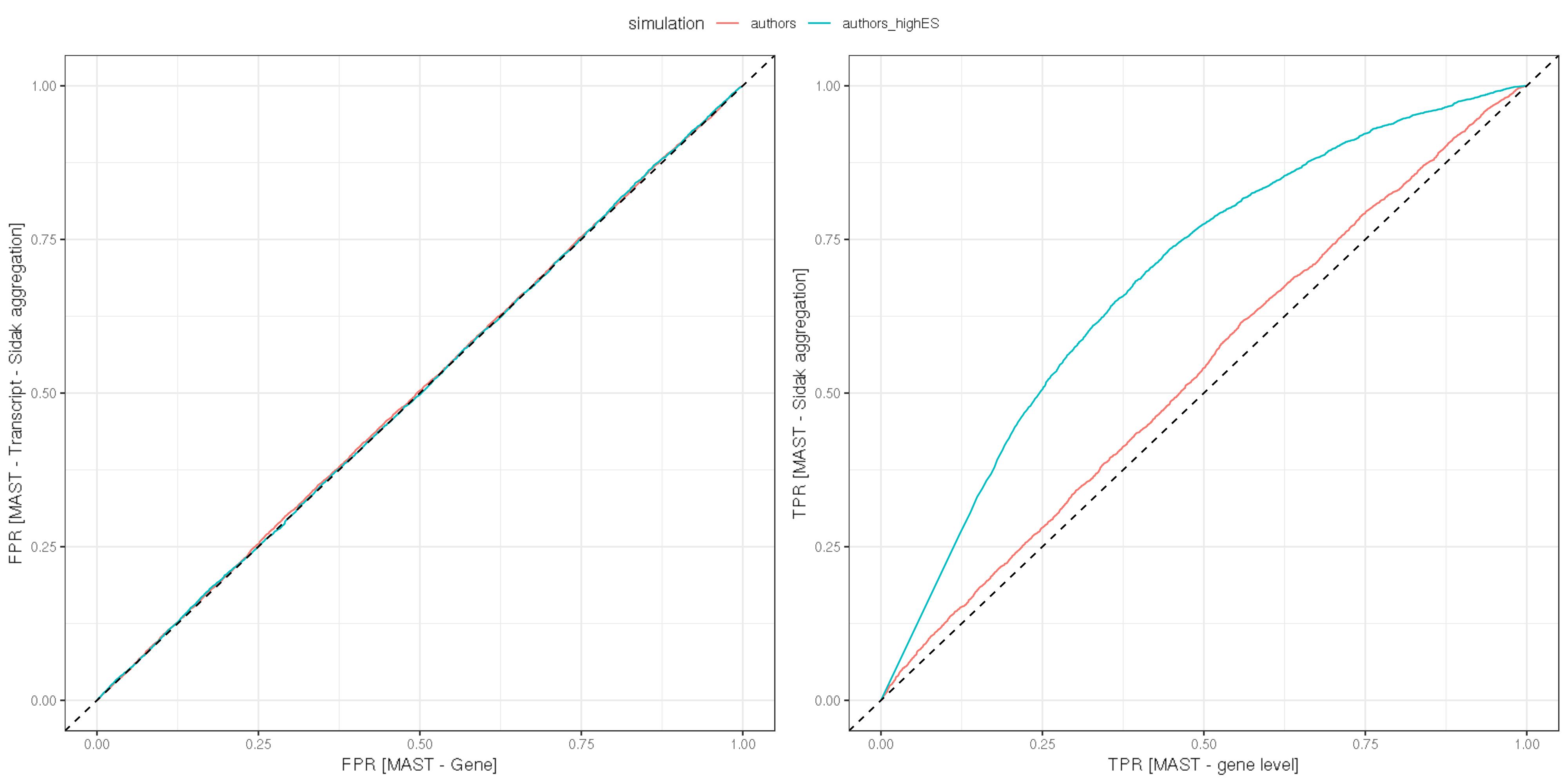

### Supplementary Figure 3

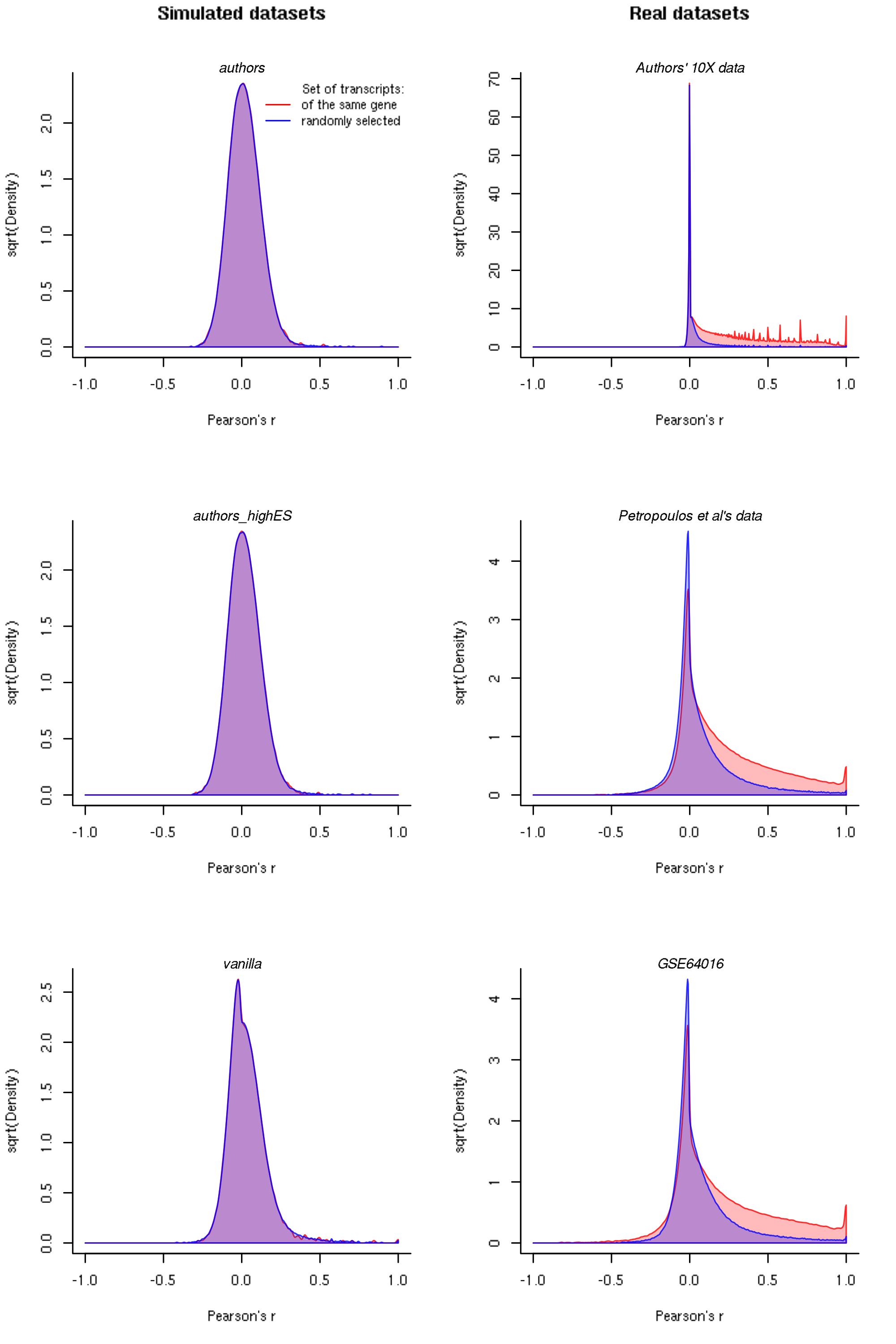
